# Default Feature Representations of the Cognitive Map

**DOI:** 10.64898/2026.05.11.724436

**Authors:** Armin Bazarjani, Payam Piray

## Abstract

Cognitive maps allow animals and artificial agents to predict where they are likely to go next, and computational models show how such predictive maps guide planning. Environments, however, rarely stay fixed. When the goal changes, a door closes, or a barrier appears, the predictive map needs to be updated. However, given that predictive maps look far ahead, a local change in the environment potentially influences what is predicted from many other states, including distant ones, even though the agent experiences it only locally. Here, we propose Default Feature Representations (DFR) to address this problem. Our solution separates a predictive map into a fixed set of features shared across environments and an operator that carries the current environment. We show that the operator has a model-based closed form when the structural change is known (DFR-MB), and we present a temporal-difference rule for learning it from sampled transitions (DFR-TD) and prove that it converges to the same solution. Using samples alone, DFR-TD efficiently learns the perturbed map, plans about as well as the model-based solution, and outperforms successor-representation baselines on replanning. The model also reproduces the local remapping of grid cells under environmental change, mirroring the stability of entorhinal grid fields across environments.^1^

## 1 Introduction

Animals navigate dynamic environments in which familiar paths are suddenly blocked, rooms are merged, and spatial boundaries shift. Adapting to such changes is a central function of the cognitive map [1–4]. Computational accounts of the cognitive map for planning fall broadly into two classes, each making a different tradeoff between flexibility and efficiency. The first class, relational-graph models [5–7] represent the environment as a graph of states and their connections, supporting strong generalization across tasks but requiring iterative computation at decision time to plan a route. The second class, predictive map models [8–14], instead cache long-range predictive structure, enabling efficient planning at decision time at the cost of needing updates when the environment changes. This second class, which includes the successor representation (SR) [8], has become a standard computational account of the cognitive map, and it is the class we focus on here. The challenge in this second class is to retain flexibility by reusing prior representations, so that a change does not require recomputation of the entire map.

One way to retain flexibility while reusing prior representations is to factorize the map into components that can be combined or updated as the environment changes. Prior featurized work has primarily applied this idea to the *reward* side of the problem: successor features [15–18] separate state features from a reward vector to enable transfer when reward changes. End-to-end factorizations of the successor measure, such as the forward-backward representation [19], contrastive RL [20], and the *γ*-model [21], likewise target reward transfer or goal-conditioned learning. When the environment itself changes, however, as when a barrier appears, a door closes, or a goal moves to a new location, these representations must be relearned from samples, either directly or together with a learned feature-space transition operator Lehnert and Littman [22], even when only a small part of the environment has changed.

A complementary line of work has approached environmental change through the structure of the map itself. Previous work [23] introduced the default representation (DR), a predictive map defined with respect to a default policy, and showed that when transitions change in a small region of the environment, the resulting change in the predictive map has low-rank structure and admits a closed-form correction expressed in terms of the original map and the local change. This idea was later extended [24] to compositional construction of cognitive maps, learning object-level predictive representations that combine to produce maps for environments with multiple objects, with applications to neural data on entorhinal cortex. Both rely on *model-based* computation: the update is derived analytically from known structural changes (a barrier, an object). An animal, however, is not told how its environment has changed. What has been missing is a model-free route to this correction, a learning rule that recovers it from local sampled experience.

In this paper we provide one. We show that the DR of an arbitrary environment can be expressed as *D* = *F M F*^*⊤*^, where *F* is a fixed feature basis derived from a symmetric base environment (such as an open-space random walk), whose DR eigenvectors have been proposed to underlie grid-cell firing [9, 23], and *M* is a change operator specific to the environment. Importantly, the basis *F* is computed once and reused across environments; *M* is what changes when the environment changes. We call this factorization *Default Feature Representations* (DFR). When the structural change is known, *M* has a closed form (DFR-MB). We also give a model-free algorithm, DFR-TD, that learns *M* from sampled transitions via a one-line semi-gradient TD update, with provable convergence to the solution under standard assumptions [25, 26]. DFR-TD thus recovers a model-based solution from model-free experience, a correspondence that, to our knowledge, has no counterpart in the factorized methods above. We further show that learning *M* from samples requires far less experience than alternative methods. The reason is that local environmental changes induce *low-rank* changes in the predictive map, and the DR factorization exposes this structure in a form that can be learned efficiently.

We validate DFR on four settings. In a *reconstruction* task, the learned *FMF* ^*⊤*^ recovers the post-change DR from sampled transitions alone after a barrier is introduced into an open field. In *replanning* tasks, including open field environments with horizontal and L-shaped barriers and four-room environments with moved goals or inserted walls, DFR-TD substantially outperforms two SR baselines, achieving the cost of the model-based optimum, while state-indexed methods remain near random-policy performance. In a *composition* task, corrections learned separately for two barriers combine without further training into an operator that achieves the model-based solution for the two-barrier environment. We further show that using targeted sampling around the barrier yields fast corrections on timescales more similar to those observed in rodents and humans. Finally, in a *remapping* setting, DFR accounts for the local remapping of grid fields under environmental change. We illustrate this by simulating the remapping experiment of Wernle et al. [27]. Together, these results show that the model-based correction to a predictive cognitive map can be learned efficiently in a model-free way from local sampled experience and entirely in the feature space, suggesting that at least part of this efficiency derives from the low-rank structure that the factorization exposes.

## 2 Default Feature Representation

### 2.1 Background and Notation

We consider a finite Markov decision process with *N* states, transition matrix *T* ∈ ℝ^*N ×N*^ under a default policy (we take this to be a uniform random walk), and per-state cost vector *c* ∈ ℝ^*N*^. The default representation [23] is

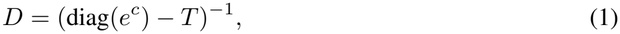

a matrix encoding long-range, cost-discounted state occupancies under the default policy. *D* supports flexible value computation: given a goal state and reward vector *r*, the optimal value function under the linear-RL framework is recovered by a single matrix-vector product involving *D* [28, 29, 23].

When the transition matrix changes from *T* to *T*_new_, for example when a barrier is introduced, the default representation changes correspondingly to

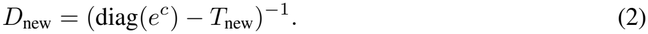

The challenge addressed in this paper is to recover *D*_new_ from sampled transitions in the new environment.

Local changes to the transition structure induce a structured, low-rank correction to the DR [23]. Suppose the change from *T* to *T*_new_ affects transitions at *B* states. Define the change *T*_new_ −*T* = *CR*^*⊤*^, where *C* ∈ℝ^*N ×B*^ is a binary indicator matrix selecting the affected states and *R*∈ ℝ^*N ×B*^ encodes the magnitude of the change at each affected state. Applying the Woodbury matrix identity [30, 31] to the definition of *D*_new_ yields

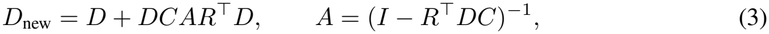

where *A* ∈ ℝ^*B×B*^. The update has rank at most *B*: if only a few transitions change, the change in *D* has correspondingly low rank.

### 2.2 Feature-Space Parameterization

Now suppose *D* is symmetric, with eigendecomposition *D* = *V* Λ*V* ^*⊤*^, and define the feature matrix *F* = *V* Λ^1*/*2^, so that *D* = *FF* ^*⊤*^. Substituting this into (3) yields

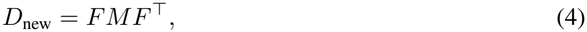

where *M* ∈ ℝ^*K×K*^ is an operator given by

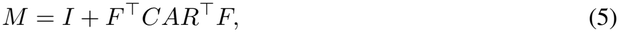

with *K* the number of features. Equation (4) is the form of the DR we work with throughout this paper. We refer to *M* as the *default feature operator*, and to the resulting parameterization *D*_new_ = *FMF* ^*⊤*^ as the *Default Feature Representation* (DFR). Intuitively, *M* re-weights and mixes the fixed feature basis to reproduce the current environment’s predictive map. We propose that this correction sits in a fast, plastic mapping between grid-cell features and hippocampal predictive codes.

The basis *F* is fixed: it depends on the base environment from which *D* was computed, not on the new environment. What changes when the environment changes is the operator *M*. Equation (5) provides a model-based expression for *M* in terms of the structural change between *T* and *T*_new_. The remainder of this section develops a model-free, sample-based learning rule for *M* that does not require knowledge of this change.

### 2.3 Bellman Equation in Feature Space

We define the rescaled feature

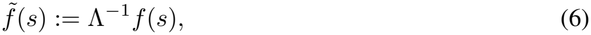

obtained by dividing each component of *f* (*s*) by the corresponding eigenvalue. Note that Λ is fully determined by *F* : the eigenvalues *λ*_*k*_ are the squared column norms of *F*. The rescaled feature can therefore be computed directly from *F* at runtime, with no auxiliary parameters.

The DR satisfies a row-wise Bellman equation that follows directly from its definition. For each state

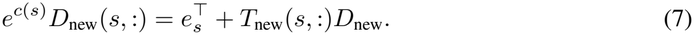

Substituting the parameterization *D*_new_ = *FMF* ^*⊤*^,

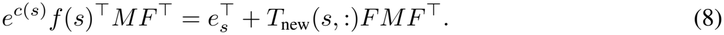

We right-multiply by the pseudoinverse (*F* ^*⊤*^)^+^ = *F* Λ^*−*1^, which exists because the columns of *F* are linearly independent (and satisfies *F* ^*⊤*^ · (*F* ^*⊤*^)^+^ = *I*_*K*_):

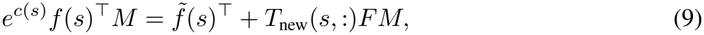

using 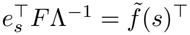. The expectation 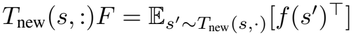 can be replaced by a sample *s*^*′*^ ~ *T*_new_(*s*, ·), yielding

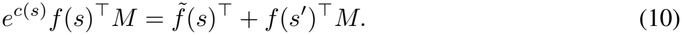

Equation (10) is exact when *K* = *N*. When *K < N*, it defines a projected Bellman equation whose solution is the best Bellman-consistent operator *M* in the basis *F*. Equation (10) is the central object of this section: it characterizes *M* in terms of quantities accessible from local experience, the features at the current state and a sampled successor, the rescaled features at the current state, and the local cost.

### 2.4 TD Learning Rule

#### Learning rule

Equation (10) suggests learning *M* by minimizing the squared Bellman residual on sampled transitions (*s, s*^*′*^). The corresponding semi-gradient TD update, treating the target as constant in *M* as is standard in TD methods [32], is

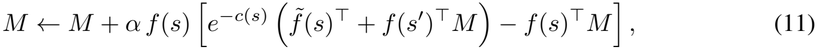

where *α >* 0 is a step size.

The update operates entirely in feature space. The state *s* enters only through *f* (*s*) and 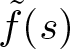, the successor *s*^*′*^ enters only through *f* (*s*^*′*^). No state-indexed quantity, such as a row of *D* or an entry of *T*, is constructed at any point in the algorithm. This is the central feature of DFR as a learning algorithm: the cognitive map is learned in the feature space defined by *F*, not in the state space defined by the environment.

Unlike the model-based form in Equation (5), the TD rule requires no knowledge of the structural change between *T* and *T*_new_. The agent observes only (*s, s*^*′*^) pairs sampled from the new environment, and updates *M* accordingly. The model-based and model-free routes converge to the same operator: the fixed point of the TD update recovers the *M* given by (5) when *K* = *N*, as we show below.

#### Convergence

The update (11) decomposes column-wise into *K* independent linear TD(0) problems, so under standard step-size conditions the iterates converge with probability 1 to a unique fixed point *M* ^*\**^. We state this formally as Theorem 1 and prove it in Appendix A. The fixed point has a clean characterization. When *K* = *N, M* ^*\**^ satisfies *FM* ^*\**^*F* ^*⊤*^ = *D*_new_ exactly, recovering the model-based expression in (5). When *K < N, M* ^*\**^ is the best Bellman-projected approximation of the true operator in the basis *F*, the standard characterization of TD(0) under linear function approximation [25, 33].

### 2.5 Choice of Feature Basis

The algorithm and convergence result make no assumptions about how *F* is obtained beyond the requirement that *D*_base_ = *FF* ^*⊤*^ for some symmetric base environment. In our experiments, the base environment is an open-space random walk, and *F* is computed once from the eigendecomposition of *D*_base_ and reused across all subsequent environments. The use of graph-operator eigenvectors as a feature basis for value approximation has a long history in reinforcement learning, originating with proto-value functions [34, 35], which use eigenvectors of the graph Laplacian as task-independent basis functions. The eigenvectors of the DR can be seen as a biologically motivated generalization of this idea. These eigenvectors have been independently connected to grid-cell response fields in entorhinal cortex [9, 23], providing a joint computational and biological motivation for this choice. The algorithmic and theoretical results, however, apply to any base environment for which *D*_base_ is symmetric.

## 3 Experiments

We evaluate six claims in turn. We measure planning performance under barrier insertion in an open field (Section 3.1), comparing our approach to another classical model-based method, and under two replanning tasks (goal- and transition-revaluation) in a structured task graph (Section 3.2). We then show that corrections learned independently for separate barriers compose without further training (Section 3.3), that learning is fast because of the basis’s alignment (Section 3.4), and that the model continues to learn when the feature basis is incomplete (Section 3.5). Finally, we show that the learned map reproduces the local grid remapping reported by Wernle et al. [27] (Section 3.6).

### Methods compared

We compare two forms of DFR against two successor representation (SR) baselines. DFR-MB is the model-based form, in which *M* is computed in closed form from the structural change between *T*_old_ and *T*_new_ via Equation (5); this requires access to *T*_new_ and serves as the planning ceiling. DFR-TD is the model-free form (Equation (11)), which learns *M* from sampled transitions in the new environment, without access to *T*_new_, and is the central contribution of this paper. SR-TD is the classical TD-learning approach to the SR [8, 11]. SR-SF is the successor-features formulation [16] adapted to learn the SR via an *N* × *K* object parameterized in a feature basis; for fair comparison, we use the same basis *F* as DFR-TD. Both baselines use the TD update Ψ(*s*) ← Ψ(*s*)+*α x*(*s*)+*γ*Ψ(*s*^*′*^)−Ψ(*s*), with *x*(*s*) = *e*_*s*_ for SR-TD (Ψ ∈ ℝ^*N ×N*^) and *x*(*s*) = *f* (*s*) for SR-SF (Ψ ∈ℝ^*N ×K*^), under the same sampling, learning rate, discount, and sample budget as DFR-TD (Appendix C). We use *c*(*s*) = 0.1 for all non-terminal states and *c*(*s*) = 0 for the terminal goal state. Planning performance is reported as cumulative cost, so higher values (closer to zero) indicate better planning. For all comparisons we assume a full feature basis (*K* = *N*), with which we found every model to perform best (see Section 3.5 for how DFR-TD handles an incomplete basis, and Appendix B.4 for low-rank versions of the baselines).

### Evaluation

Planning performance is measured as the mean accumulated cost across a fixed set of evaluation rollouts. Each representation yields a value function via a single matrix-vector product, from which we roll out a softmax policy until the goal is reached or 1k steps elapse; all methods share the same planner, temperature, step cap, and evaluation pairs, so only the representations differ (Appendix C). For each task, start and goal regions lie on opposite sides of the relevant perturbation (a barrier or a moved goal), so that solving the task requires the agent to route around it. Every experiment is run across 5 random seeds. Within a seed, open field performance is the mean over 200 fixed (start, goal) pairs, and four-room performance is the mean over 20 rollouts. Learning curves show the mean with a shaded 95% confidence interval; tables report the mean with SEM in parentheses.

### 3.1 Planning around inserted barriers in an open field

Before evaluating planning, we verify that DFR-TD recovers the perturbed predictive map purely from on-policy experience in the new environment, with no access to its transition matrix. Inserting a horizontal barrier into an open field and training *M* using Equation (11), we find that the median reconstruction error |*FMF* ^*⊤*^ −*FM*_true_*F* ^*⊤*^| decreases steadily with training across feature dimensions *K* ∈ {10, 25, 50, 75, 100}, with larger *K* generally reaching lower asymptotic error (Appendix B.1). The TD rule recovers the model-based solution from samples alone, as Theorem 1 guarantees.

We next ask whether the learned *M* can support planning, and how it compares to baselines. We use two open field configurations differing in barrier geometry: a horizontal barrier (Figure 1a) and an L-shaped barrier (Figure 1b). For each condition we insert the aforementioned barrier into the environment, changing the transition structure *T*_old_ → *T*_new_. Critically, none of the agents, apart from the optimal DFR-MB agent, had direct access to the transition structure and they all had to learn the changes online through direct experience.

**Figure 1:**
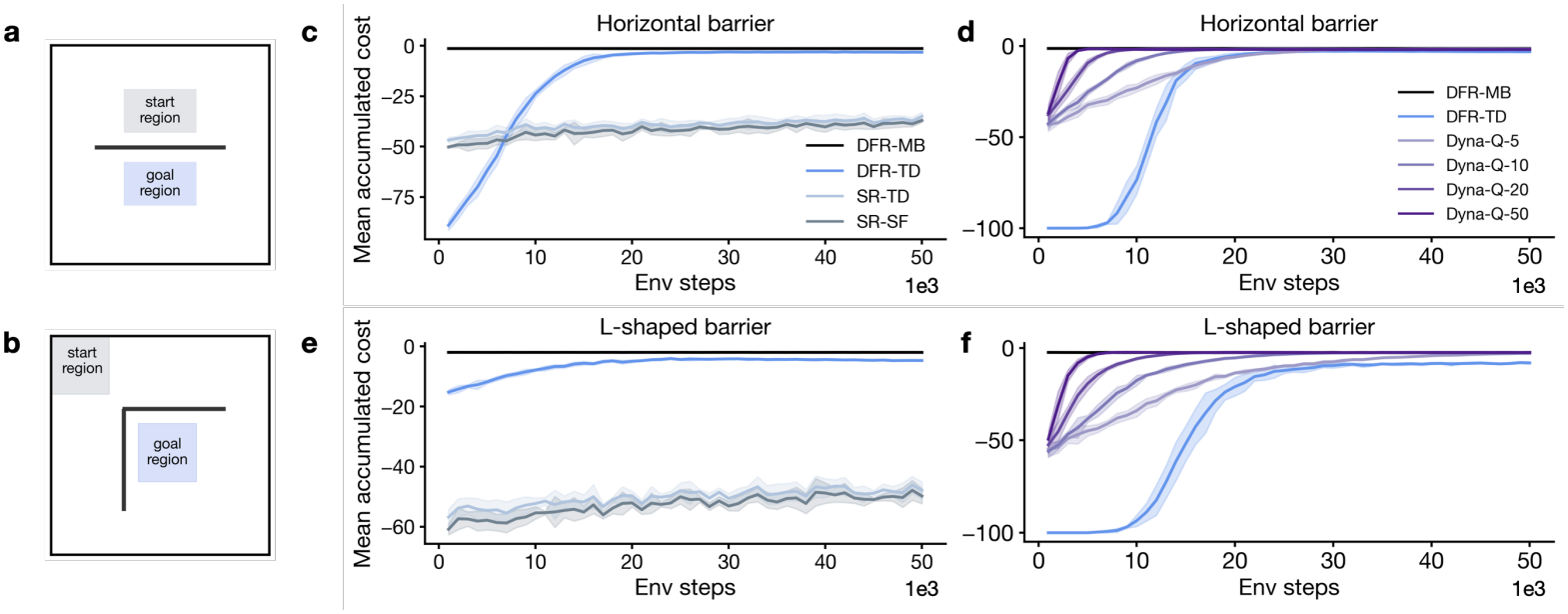
Planning under barrier insertion in an open field. **(a, b)** Open field environments with a horizontal **(a)** and L-shaped **(b)** barrier; gray and blue squares mark evaluation start and goal regions, respectively. **(c-f)** Mean accumulated cost across evaluation rollouts as a function of environment steps. DFR-TD (blue, ours) reaches the model-based DFR-MB ceiling (black), while SR-TD (light blue) and SR-SF (dark gray) plateau below. Dyna-Q with varying numbers of rollouts (shades of purple) also converges to the model-based ceiling. Shaded regions denote 95% CI.

The goal of this setting was to evaluate how well each model can update its representation to accommodate the barrier. A perfectly updated representation will reflect the barrier’s geometry and thus yield fewer steps from start to goal. For both conditions (horizontal and L-shaped), evaluation start and goal regions lie on opposite sides of the barrier, we sample 200 different (start, goal) pairs and intermittently evaluate the model’s ability to plan to the goal state as it updates its representation. Both configurations exhibit the same pattern (Figure 1c, e): DFR-TD attains planning performance similar to the optimal DFR-MB model while neither SR-TD nor SR-SF are able to update their representations in any meaningful manner. Note that the efficiency arises not from compressing the map but from the structure of the update: a local change induces a low-rank change in the operator. The DR factorization exposes this in a form that TD can learn quickly. A basis comparison at fixed *K* = *N* (Section 3.4) confirms this: bases from related environments perform comparably to the open field basis, while a random basis with matched spectrum and dimension is far slower.

Next, we evaluate our model against Dyna-Q [36, 37], an integrated planning algorithm that builds an explicit model of the environment from sampled transitions and interleaves real experience with simulated backups (Figure 1d, f). Dyna-Q achieves final performance comparable to DFR-MB after sufficient samples, as expected for a method with access to a learned model of *T*_new_. DFR-TD achieves similar performance while learning a low-rank correction to an existing map from samples, rather than learning a model and planning with it.

### 3.2 Replanning under goal and transition changes in a structured environment

In this set of experiments we probe how well the model handles different replanning paradigms, a central capacity that any model of the cognitive map should support [1]. For our simulations we used the familiar four-room environment as it provides a structured counterpart to the open field.

#### Goal revaluation

To test replanning in the face of a changing goal location we initially placed the goal at *r*_1_ (top-right corner of room TR; Figure 2a). We then moved the goal to one of the eight other corner locations *r*_2_, …, *r*_9_. We separate the evaluations into two conditions: *same room*, in which the new goal lies in the same room as the original (*r*_2_, *r*_3_), and *different room*, where it has been moved into a different room than the original (*r*_4_, …, *r*_9_). Although a goal change might appear to alter only the reward function, goals also act as absorbing terminal states; consequently, moving the goal modifies *T* at both the previous and new goal locations, and the predictive map must be updated. As shown in Figure 2b, DFR-TD again quickly reaches the model-based (DFR-MB) ceiling while both SR-TD and SR-SF trail behind. Unsurprisingly we see that when the goal change is in the same room, the model needs very few steps to update its representation as the previous one it learned is permissible for the majority of the required planning.

**Figure 2:**
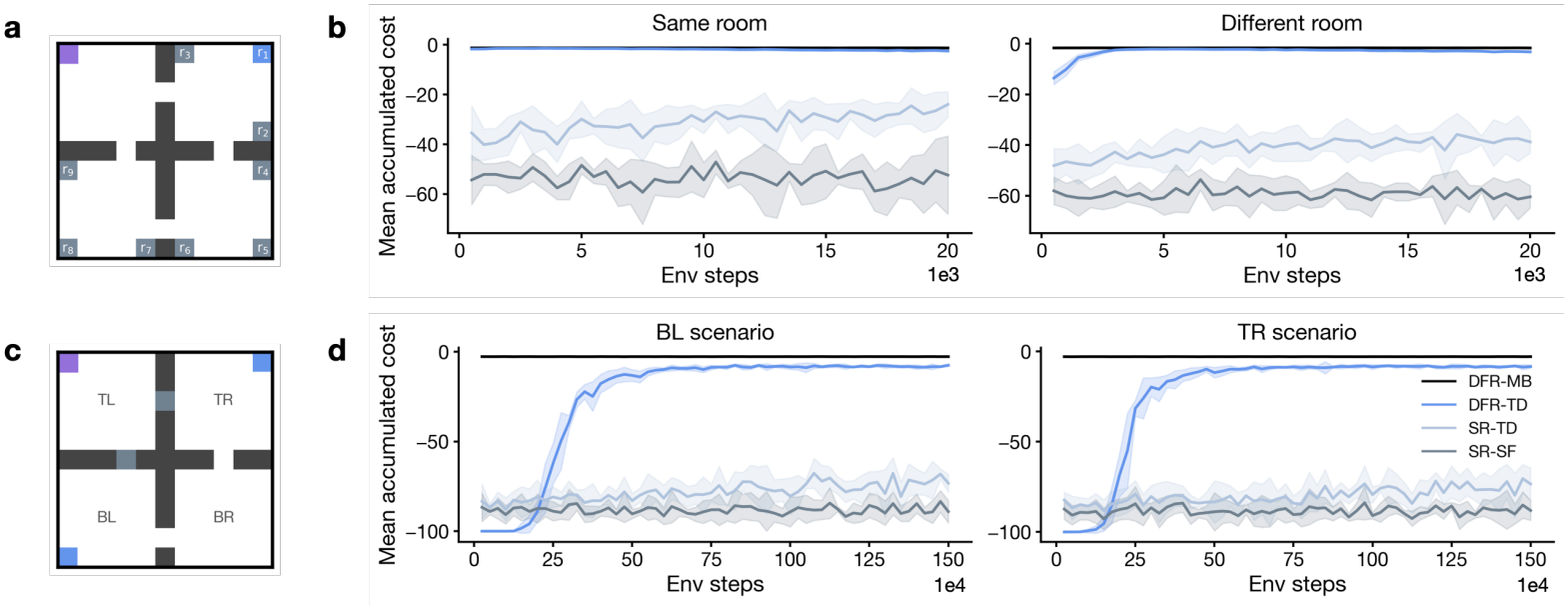
Replanning in the four-room environment. **(a)** Goal-revaluation task: a goal initially placed at *r*_1_ is moved to one of eight corner positions *r*_2_, …, *r*_9_; the agent always starts in the top-left corner of room TL (purple square). Goals at *r*_2_ and *r*_3_ constitute the *same room* condition; goals at *r*_4_, …, *r*_9_ constitute the *different room* condition. **(b)** Mean accumulated cost in the same-room (left) and different-room (right) conditions. **(c)** Transition-revaluation task: starting from a learned goal-conditioned map (goal in BL or TR, blue squares), a barrier (gray bar) blocks the direct doorway from TL into the goal room. **(d)** Mean accumulated cost in the BL (left) and TR (right) scenarios. Across both regimes, DFR-TD reaches the model-based ceiling, while SR baselines remain trapped near the pre-perturbation cost. Shaded regions denote 95% CI.

#### Transition revaluation (detour)

A classic test of cognitive flexibility is the detour task [1]: how does an agent respond when a previously learned pathway is suddenly blocked? A truly “cognitive”agent, like its human counterparts, should be able to quickly accommodate the change in the environment and find a new path to its destination.

We simulate this by starting from a learned map for a goal in room BL or TR, as indicated by the blue squares in Figure 2c. We then insert a barrier blocking the direct doorway from the start room (TL) into the goal room (Figure 2c). The agent, fixed to start in the top-left corner of TL, must learn to route through an alternative doorway to reach the goal. This setting serves as a strict generalization of the Tolman detour task. DFR-TD shows this flexibility, converging within 5–10 units of the model-based ceiling in both BL and TR scenarios (Figure 2d), while the SR baselines remain near their pre-perturbation cost throughout.

Together, the open field and four-room experiments show that our method matches model-based planning and replanning performance, without requiring *T*_new_, across all four perturbation types tested: horizontal barrier, L-shaped barrier, goal revaluation, and doorway closure. DFR-TD is also faster than its other predictive map counterparts in replanning capabilities. Final performance for every model across both environments is tabulated in Appendix B.2.

### 3.3 Composing learning of two barriers for single-shot planning

We wanted to test if the corrections learned through DFR-TD compose as composition yields a testable circuit-level claim: corrections acquired for individual objects should combine additively when their effects are roughly independent. In a new experiment two barriers are introduced and their corrections *M*_1_ and *M*_2_ learned separately (Figure 7). We then compose the two independently learned corrections into the compositional operator *M* = *M*_1_ + *M*_2_ − *I*.

The model is never trained in the two-barrier environment: *M*_1_ and *M*_2_ are learned independently and composed at test. The zero-shot composed operator approaches the model-based oracle, whereas either correction alone does not (Appendix B.6).

In that experiment, each correction was learned for 100k steps from a random walk across the whole environment at a learning rate of 0.01 (Appendix C). That regime is a poor model of how an animal would acquire the correction. An animal that encounters a new barrier would likely investigate it locally and update its representation quickly. We therefore reran the experiment with 90% of transitions sampled from states adjacent to each barrier (versus ~9% under the random walk) and a higher learning rate. Table 1 reports the mean number of steps to goal of the composed operator across 200 (start, goal) pairs: roughly 800 local transitions per barrier suffice, within the range of exposure an animal would accumulate while investigating a novel barrier.

**Table 1:** Fast learning of composition. Mean steps to goal of the composed operator vs. local transitions per barrier.

| Steps | lr = 0.1 | lr = 0.2 |
|---|---|---|
| 200 | 346.1<sup>(±56.5)</sup> | 182.3<sup>(±38.7)</sup> |
| 400 | 169.3<sup>(±43.3)</sup> | 78.3<sup>(±20.6)</sup> |
| 600 | 64.9<sup>(±18.2)</sup> | 31.3<sup>(±4.4)</sup> |
| 800 | 25.8<sup>(±1.7)</sup> | 25.9<sup>(±1.6)</sup> |

### 3.4 Why learning is fast: a basis comparison

The advantage of DFR-TD persists at *K* = *N*, where the representation is full rank and no dimensionality reduction takes place. What accounts for it there? We ran a comparison in the open field with a horizontal barrier and in the four-room environment with a barrier insertion, varying only the basis *F* used to construct the feature representation.

A basis from a different but similar environment (the L-barrier map in the open field and the four-room map with the other door blocked) converges to the same performance as the open field basis on both tasks, as does the generic open field basis in the four-room (Figure 3). The method therefore works across a family of structurally reasonable bases, not a single privileged one. A random orthonormal basis with matched spectrum and identical dimension, by contrast, is dramatically slower on the barrier task and fails the four-room task, despite identical size and identical asymptotic solution. A poorly chosen *F* hurts only when it carries no relevant structure at all; across the bases an agent would plausibly hold, performance is robust. We reason that what matters is that *F* is a DR basis, not which environment’s.

**Figure 3:**
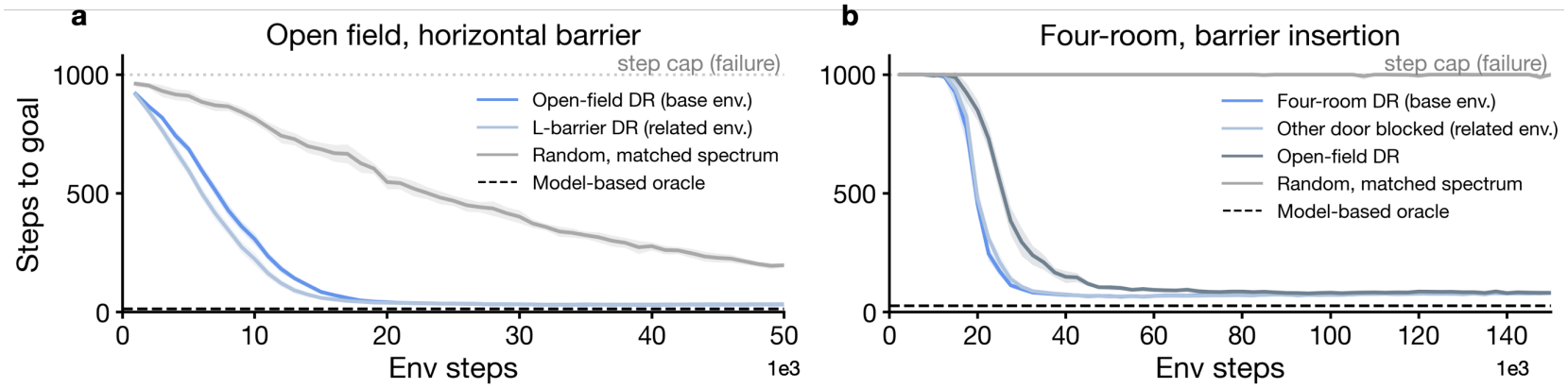
Basis comparison. (**a**) Open field with a horizontal barrier. (**b**) Four-room with a barrier insertion. Every basis has the same size, the same TD fixed point, and the same starting map; only its alignment differs. DR bases from the base or a related environment learn at the same rate, while a random basis with matched spectrum is far slower (**a**) or never reaches the goal within the step cap (**b**). Dashed: model-based oracle. Shaded regions denote 95% CI.

**Figure 4:**
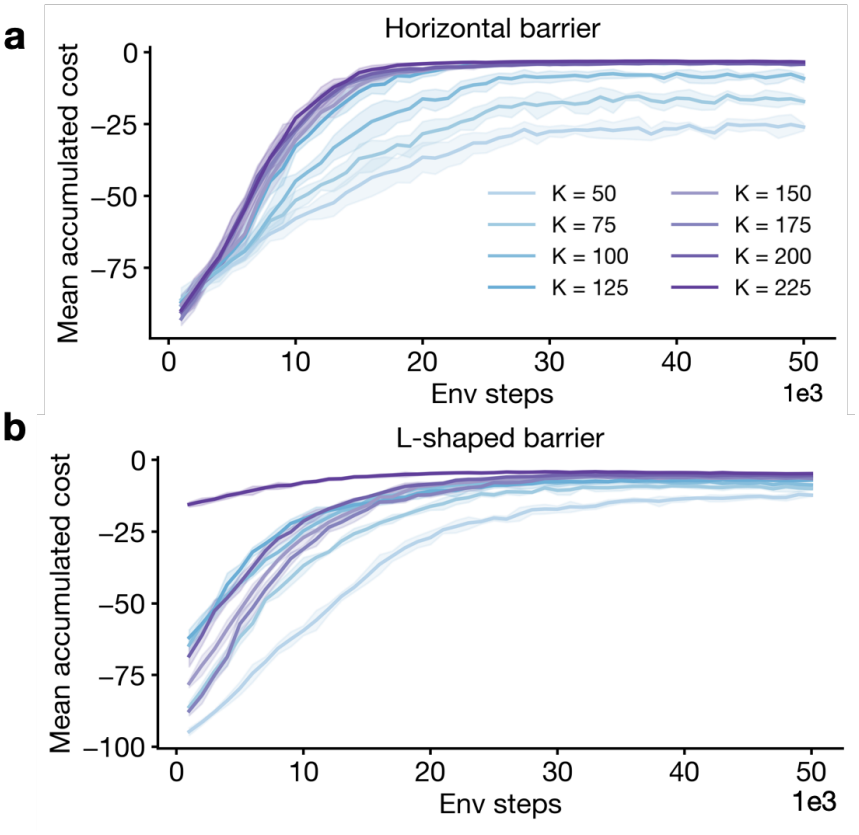
Planning performance of DFR-TD with a reduced feature space. **(a, b)** Planning results in open field environment with a horizontal **(a)** and L-shaped **(b)** barrier. As the feature space becomes richer (higher *K*) the model learns quicker and converges to the optimal solution. Shaded regions denote 95% CI.

The DR basis is tied to the environment’s transition structure through Equation (1) and Equation (4). Because *F* is constructed from *D*, the base environment corresponds to *M* = *I*, and a local change adds a correction of rank at most *B* on top of it, even though its dimension is the same as *D*. That correction is what the learner acquires. For a basis not derived from *D*, the operator to be learned is full rank. The rank of the learning problem is therefore set by the basis together with the environmental change, which is why a random basis of identical size and matched spectrum is slow. Read this way, the comparison is a robustness result: the advantage holds across DR bases, as long as the basis stays within that family. The comparison in Figure 3 shows that the advantage is relatively robust across bases drawn from the same DR family (full results in Appendix B.7) (full results in Appendix B.7).

### 3.5 Can DFR-TD accommodate an incomplete feature space

One of the final tests we performed on the model was to see how well the approach could accommodate a reduced feature space. That is, can it handle an “incomplete” feature space, as is often the case in the real world? This is not a claim that DFR benefits from a smaller basis, but a test of whether it survives an imperfect one, the realistic case for any basis derived from finite experience or instantiated in a biological system. In the previous experiments, we assumed a complete feature space and allowed our model to have access to all of the eigenvectors provided by the spectral decomposition. We now test the model by only allowing it to have access to the top-*K* eigen-vectors of *D*. As expected, larger *K* yields faster learning and better final performance. More notably, DFR-TD continues to plan rea-sonably well even when *K* is substantially smaller than *N*, despite the basis no longer being able to exactly reconstruct *D*. This robustness likely reflects a property of the TD objective: because it minimizes a projected Bellman error in whatever basis is supplied, it can tolerate an imperfect decomposition that the closed-form update cannot. DFR-MB, which requires exact decomposition, fails sharply under the same truncation (Appendix B.3, Table 4). The contrast is informative: DFR-TD degrades gracefully, DFR-MB does not.

This raises a practical question: if the basis may be incomplete, can the agent tell? Appendix B.5 shows that the TD error of Equation (11), averaged over the states the agent visits, decreases monotonically with K and tracks the reconstruction error against the true post-change map, making it usable as an internal signal of basis adequacy that requires no access to that map, and as a trigger for growing *K* online.

### 3.6 The learned map reproduces local grid remapping

Finally, we ask whether a learned transformation, *M*, exhibits the spatial structure observed in grid cells in the medial entorhinal cortex (MEC) under environmental change. To test this, we replicate the merging experiment of Wernle et al. [27]. In their setup, two adjacent rectangular compartments, separated by a central partition during training, are merged at test time when the partition is removed. The experimental finding is that grid cell responses remain stable away from the former partition but reorganize locally near it, producing a characteristic low-correlation band in the population sliding-correlation map.

As in prior accounts of grid cells as a low-dimensional spectral representation of the cognitive map [9, 24], we relate grid cells to a spectral basis derived from the predictive map. We compute *F* from the transition structure of the divided environment that contains the partition between the two compartments. Upon removal of the partition, the transition structure changes; applying the TD rule of Equation (11) yields an operator *M* that transforms the basis into *F*_new_ = *FM*. We compute the sliding spatial correlation between *F* and *F*_new_ across the population, following the analysis of Wernle et al. [27]. The model reproduces the key experimental finding: correlation is highest in regions distant from where the partition once stood (Figure 5).

**Figure 5:**
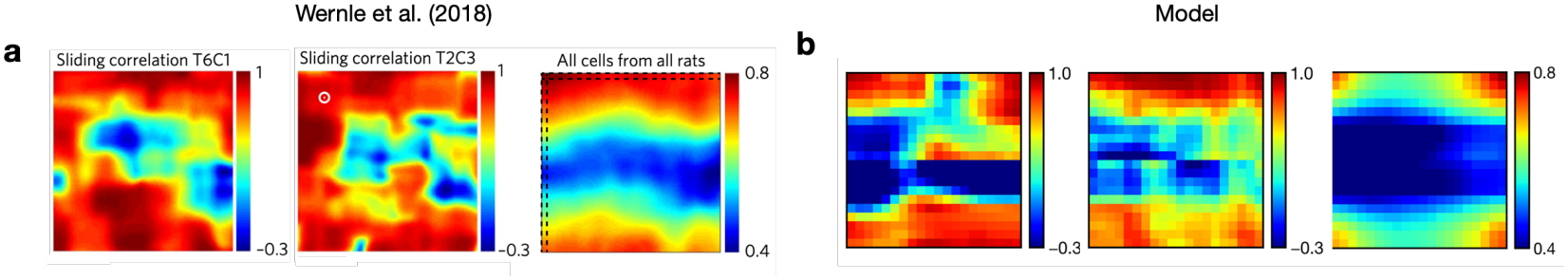
The learned map reproduces local grid remapping. (**a**) Data from Wernle et al. [27]: sliding spatial correlation for two example cells (left, center) and the population average over n = 128 cells (right). (**b**) Model: the same analysis for two of the model’s cells (left, center) and the population average over all of the model’s cells (right).

Crucially, this prediction is recovered without any access to *T*_new_; it follows from the transformation imposed by the learned operator *M* alone. This extends the analytical model-based result of prior work [24] to the experientially-grounded case in which the perturbation must be discovered through interaction.

## 4 Discussion

We introduced Default Feature Representations (DFR), a framework for learning and updating a featurized cognitive map from sampled experience. The core insight is that the DR of any environment can be expressed as *FMF* ^*⊤*^, where *F* is a fixed spectral basis derived once from a base environment and *M* is a change operator that absorbs all environment-specific structure. The model-free temporal-difference rule learns *M* from transitions in the new environment without requiring access to the transition matrix. Across reconstruction, planning, composition, and neural-replication tasks, DFRTD matches the performance of the model-based analytic update (DFR-MB) while operating entirely from sampled experience, and outperforms SR-TD and SR-SF across all conditions tested. Successor features already allow the post-change map to be learned without a model; what DFR adds is that it learns a correction to the existing map entirely in the feature space, rather than relearning a state-indexed map, and this is what makes it fast. DFR thereby carries the analytic account of how a predictive map should change [23, 24] into the setting an animal actually faces, where the change must be discovered through interaction, using only local experience and a one-line TD update.

The DFR factorization *D* = *FMF* ^*⊤*^ imposes a natural two-timescale separation on how the cognitive map changes. The feature basis *F*, the eigenvectors of the open-space DR, which we identify with grid cells in the MEC [23], is stable across environments and changes only on the timescale of the animal’s overall spatial experience. The operator *M*, by contrast, is environment-specific and updates quickly from local transitions when barriers are inserted or goals are moved. Under this account, the underlying coordinate system *F* is preserved across environments; what changes is the operator *M* that combines basis elements into the predictive geometry of the current environment. Empirically observed grid-cell remapping [27] then reflects the change in *M*, not a change in the basis itself. Similarly, this can be further contextualized within the broad class of models of cognition employing both a fast and slow learner [38–40].

This picture aligns with a broader hypothesis of a two-timescale separation in the entorhinal-hippocampal system. Grid cells in the MEC provide a stable, metric coordinate system [41, 42], a set of spatial features that is largely preserved and reused across environments [43], while place cells in the hippocampus exhibit environment-specific remapping [44, 45] and object-vector cells in the MEC fire at a particular distance and direction from objects, wherever they are placed [46]. The DFR framework offers a computational rationale for this division of labor: *F* is the grid-cell coordinate system, reused across environments, while *M* is the environment-specific transform that instantiates a new place-cell map without requiring the grid system to be re-acquired; corrections for separate objects then combine within *M* (Section 3.3), as in Piray and Daw [24].

## Limitations

The most obvious constraint is that *F* is fixed, derived once from an open-space random walk and reused across all tested environments. When the base geometry differs substantially from the perturbed one, the approximation *D*_new_ ≈ *FMF* ^*⊤*^ may degrade and the appropriate *F* to use is not obvious a priori. The convergence guarantee (Theorem 1) also assumes transitions are sampled from the stationary distribution of *T*_new_, whereas in practice the agent must first explore the perturbed environment, and how badly planning suffers during this transient phase depends on the exploration policy in ways we have not characterized. We have also modeled environmental change as a perturbation from a fixed baseline, which fits naturally for barrier insertion and goal relocation but may not extend cleanly to more drastic structural changes. Our comparison against low-rank successor representations (Appendix B.4) uses a single environment and does not show that DFR is the only route to a compact, adaptable map; compositionality (Section 3.3) is shown only for approximately non-interacting changes; and how to choose *K*, and when to grow it, in the truncated regime remains open (Appendix B.5). Finally, all experiments use tabular gridworlds, and whether DFR’s sample-efficiency advantage survives contact with continuous or high-dimensional state spaces remains an open question.

## Future directions

Several extensions follow naturally. Most directly, the fixed-*F* assumption could be relaxed by jointly learning *F* and *M* on two different timescales: slow updates to *F* tracking gradual changes in global spatial layout, and fast updates to *M* accommodating local perturbations. A change that persists across episodes is cheaper to amortize into the basis than to carry indefinitely in *M*, mirroring the reorganization of grid maps under persistent restructuring [47]. This proposal is conceptual rather than a proven guarantee: Theorem 1 assumes a fixed *F*, so convergence for a jointly updated (*F, M*) is a well-defined next step, although the success of bases from related environments (Section 3.4) suggests *F* can be updated slowly without destabilizing the fast *M* correction. This would formalize the two-timescale hypothesis above and connect DFR to accounts of habitual behavior in which slowly updated default policies coexist with fast goal-directed planning [23, 48]. Second, Section 3.3 suggests that *M* admits compositional decomposition along the lines of Piray and Daw [24]; extending this to interacting objects is a natural next step. Finally, the connection to the entorhinal-hippocampal system raises the possibility of testing DFR’s predictions against single-unit recordings during dynamic environment changes, particularly the prediction that grid-cell remapping should be governed by a uniform linear transform *M* across the entire environment, with the magnitude of remapping peaking near the locus of structural change.

## Conclusion

The core message of this paper is that featurizing the cognitive map is not merely a computational convenience: it exposes a structural decomposition of how the map should change. The feature basis *F* captures the stable spatial geometry of the world; the operator *M* captures how that geometry is distorted by local environmental change. Updating *M* via the TD rule (Equation (11)) requires only samples in the new environment and recovers a map that matches the model-based optimum in both planning performance and neural predictions. We hope this provides a principled, biologically grounded account of how the brain updates its cognitive map from local experience.

## Funding

This work was supported by an NIMH grant (R21MH134217) and a National Science Foundation CAREER Award (2543567) to Payam Piray.

## Competing interests

The authors declare no competing interests.

## A Convergence proof

### Theorem 1.

*Suppose transitions* (*s, s*^*′*^) *are sampled with s drawn from the stationary distribution µ of T*_*new*_ *and s*^*′*^ ~ *T*_*new*_(*s*, ·). *Suppose further that the per-state cost satisfies c*(*s*)*>* 0 *for all s, the feature matrix F has full column rank, and the step sizes satisfy* Σ _*t*_ α_*t*_ = ∞ *and* 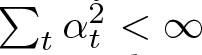.*then the iterates produced by the TD update* (11) *converge with probability 1 to a unique fixed point M* ^*\**^.

*Proof*. The TD update (11) decomposes column-wise. Writing *m*_*k*_ for the *k*-th column of *M* and *r*_*k*_(*s*) := *e*^*−c*(*s*)^*f*_*k*_(*s*)*/λ*_*k*_ for a per-column reward, the update becomes

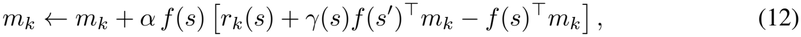

which is the standard linear TD(0) update for the value function *V*_*k*_(*s*) = *f* (*s*)^*⊤*^*m*_*k*_ in a Markov reward process with state-dependent discount *γ*(*s*) := *e*^*−c*(*s*)^, reward *r*_*k*_(*s*), and feature vector *f* (*s*).

Following *T*_new_ in the new environment induces a sampling distribution over states; we denote this distribution *µ*, the stationary distribution of *T*_new_. Taking the expectation of the update direction with respect to *µ*,

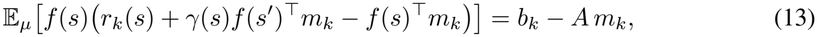

where

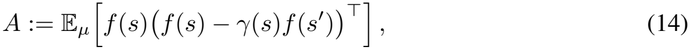

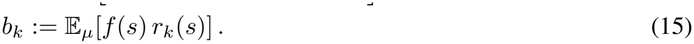

This is the same structure as the TD(0) fixed point analyzed by Tsitsiklis and Van Roy [25], with the only difference being that the discount factor *γ*(*s*) = *e*^*−c*(*s*)^ is state-dependent rather than constant. As in their analysis, convergence with probability 1 follows under standard step-size conditions whenever *A* has eigenvalues with strictly positive real parts. We verify this for our setting below, with the state-dependent *γ*(*s*) handled by a simple bounding argument.

For any *x* ∈ ℝ^*K*^, let *v* := *Fx* ∈ ℝ^*N*^. Then

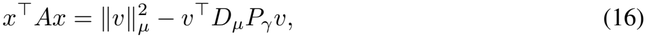

where 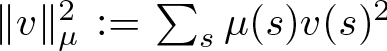, *D*_*µ*_ := diag(*µ*), and *P*_*γ*_ := diag(*γ*)*T*_new_. Cauchy–Schwarz in the *µ*-weighted inner product gives ∥*v*^*⊤*^*D*_*µ*_*P*_*γ*_*v*|≤ ∥ *v*∥ _*µ*_ ·∥*P*_*γ*_*v*∥ _*µ*_. Since *T*_new_ is row-stochastic with stationary distribution *µ*, Jensen’s inequality gives ∥*T*_new_*v* ∥_*µ*_ ≤ ∥*v*∥ _*µ*_, and combined with the diagonal scaling,

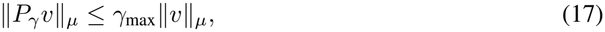

where *γ*_max_ := max_*s*_ *γ*(*s*). Therefore,

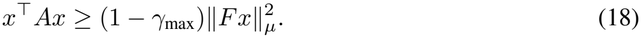

The assumption *c*(*s*) *>* 0 for all *s* gives *γ*_max_ *<* 1, and full column rank of *F* gives 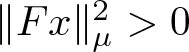 for all *x* ≠0. Hence *A* is positive definite. By Tsitsiklis and Van Roy [25], each column *m*_*k*_ converges with probability 1 to the unique fixed point 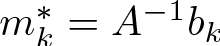. The matrix iterates converge to 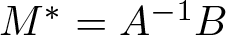 with *B* := [*b*_1_, …, *b*_*K*_].

When *K* = *N, M* ^*\**^ satisfies *FM* ^*\**^*F* ^*⊤*^ = *D*_new_ exactly, recovering the model-based expression in Equation (5). When *K < N, M* ^*\**^ is the best Bellman-projected approximation in the basis *F* [25, 33].

## Supplementary analysis

### B.1 The TD rule recovers the perturbed predictive map

The most basic claim of DFR-TD is that *M* can be learned from on-policy experience in the perturbed environment without any access to its transition matrix. We verify this directly. We insert a horizontal barrier in an open field (Figure 6a) and use the decomposition from the open field, *F*, to train *M* using Equation (11). We report the median absolute reconstruction error |*FM*_true_*F* ^*⊤*^− *FMF* ^*⊤*^|, where *M*_true_ is calculated analytically using Equation (5) (Figure 6b).

**Figure 6:**
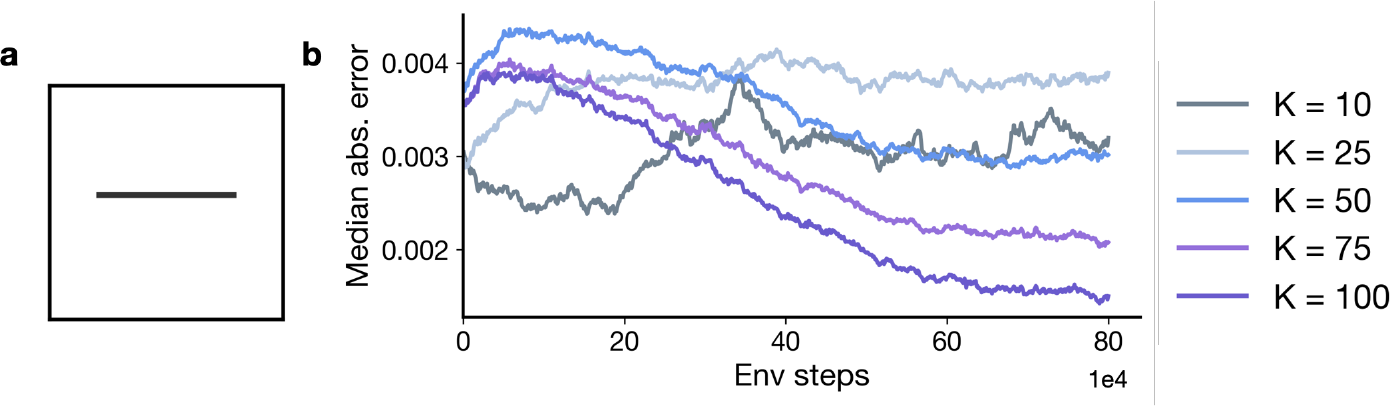
The TD rule recovers the perturbed predictive map. **(a)** Open field environment with an inserted horizontal barrier. **(b)** Median absolute reconstruction error |*FM*_true_*F* ^*⊤*^ −*FMF* ^*⊤*^ |as a function of environment steps, for feature dimensions *K* ∈ {10, 25, 50, 75, 100}. Larger *K* generally achieves lower asymptotic error.

**Figure 7:**
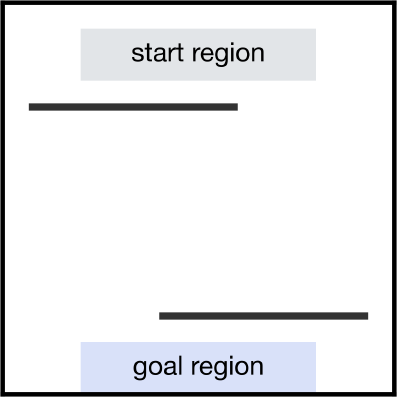
Two-barrier environment. One barrier is inserted near the start region and another near the goal region. Each barrier is learned independently of the other.

Reconstruction error decreases steadily with training across all feature dimensions *K* ∈ {10, 25, 50, 75, 100}, and generally reaches a lower asymptote at larger *K*. These results confirm that TD rule (Equation (11)) recovers the perturbed map purely from samples in the new environment.

### B.2 Final step means across models and environments

Because the cost plots can be difficult to read, we also provide the mean performance of each model we tested across both conditions in both environments. We report the final mean number of steps the model takes in each condition. Table 2 reports the final mean performance of each model across both open field environments while Table 3 reports the final mean performance of each model across both conditions of both revaluation paradigms in the four-room environment.

**Table 2:** Final performance across open field tasks.

|  | Conditions |  |
|---|---|---|
| Model name | Horizontal | L-shaped |
| DFR-MB | 13.7<sup>(±0.1)</sup> | 22.1<sup>(±1.3)</sup> |
| DFR-TD | 32.0<sup>(±0.4)</sup> | 47.2<sup>(±0.7)</sup> |
| SR-TD | 349.8<sup>(±7.4)</sup> | 478.7<sup>(±0.7)</sup> |
| SR-SF | 370.5<sup>(±3.4)</sup> | 497.9<sup>(±9.1)</sup> |
| Dyna-Q-5 | 16.5<sup>(±0.1)</sup> | 29.2<sup>(±0.4)</sup> |
| Dyna-Q-10 | 19.7<sup>(±0.1)</sup> | 24.3<sup>(±0.1)</sup> |
| Dyna-Q-20 | 19.8<sup>(±0.1)</sup> | 25.4<sup>(±0.1)</sup> |
| Dyna-Q-50 | 19.8<sup>(±0.1)</sup> | 25.4<sup>(±0.1)</sup> |

**Table 3:** Final performance across four-room tasks.

|  | Goal revaluation |  | Transition revaluation |  |
|---|---|---|---|---|
| Model name | Same room | Different room | BL scenario | TR scenario |
| DFR-MB | 12.7<sup>(±0.1)</sup> | 16.7<sup>(±0.1)</sup> | 26.7<sup>(±0.1)</sup> | 28.9<sup>(±0.1)</sup> |
| DFR-TD | 25.0<sup>(±1.3)</sup> | 32.0<sup>(±1.5)</sup> | 74.7<sup>(±1.9)</sup> | 83.8<sup>(±3.3)</sup> |
| SR-TD | 239.6<sup>(±18.6)</sup> | 388.0<sup>(±15.9)</sup> | 697.2<sup>(±45.7)</sup> | 809.3<sup>(±28.2)</sup> |
| SR-SF | 523.9<sup>(±56.6)</sup> | 604.6<sup>(±15.9)</sup> | 890.6<sup>(±22.5)</sup> | 881.2<sup>(±19.5)</sup> |

**Table 4:** Final performance with reduced feature space across open field tasks.

|  | Horizontal |  | L-shaped |  |
|---|---|---|---|---|
| Feature dim (K) | DFR-MB | DFR-TD | DFR-MB | DFR-TD |
| 50 | 738.2<sup>(±13.5)</sup> | 259.5<sup>(±4.6)</sup> | 985.2<sup>(±1.2)</sup> | 122.6<sup>(±1.7)</sup> |
| 75 | 679.7<sup>(±9.1)</sup> | 170.6<sup>(±3.7)</sup> | 966.5<sup>(±1.3)</sup> | 99.7<sup>(±1.6)</sup> |
| 100 | 695.2<sup>(±4.3)</sup> | 90.3<sup>(±5.3)</sup> | 780.0<sup>(±3.6)</sup> | 87.2<sup>(±2.5)</sup> |
| 125 | 609.1<sup>(±5.4)</sup> | 36.3<sup>(±1.1)</sup> | 925.4<sup>(±2.2)</sup> | 70.2<sup>(±0.9)</sup> |
| 150 | 710.1<sup>(±8.2)</sup> | 38.2<sup>(±0.9)</sup> | 987.0<sup>(±2.4)</sup> | 64.9<sup>(±1.9)</sup> |
| 175 | 847.3<sup>(±6.4)</sup> | 40.8<sup>(±0.8)</sup> | 991.4<sup>(±0.3)</sup> | 63.4<sup>(±0.9)</sup> |
| 200 | 734.1<sup>(±13.2)</sup> | 40.9<sup>(±0.9)</sup> | 986.9<sup>(±0.9)</sup> | 54.1<sup>(±0.8)</sup> |
| 225 | 13.8<sup>(±0.1)</sup> | 33.3<sup>(±0.8)</sup> | 18.8<sup>(±0.1)</sup> | 47.3<sup>(±0.3)</sup> |

**Table 5:** Comparison with low-rank successor representations across feature dimensions.

| *k* | SR-SF-*k* | SR-LR | DFR-TD |
|---|---|---|---|
| 50 | 709.4<sup>(±9.6)</sup> | 508.5<sup>(±3.9)</sup> | 294.4<sup>(±13.0)</sup> |
| 75 | 587.2<sup>(±7.0)</sup> | 486.4<sup>(±5.7)</sup> | 184.7<sup>(±9.5)</sup> |
| 100 | 537.5<sup>(±4.4)</sup> | 497.9<sup>(±4.8)</sup> | 97.2<sup>(±6.0)</sup> |
| 125 | 488.4<sup>(±3.5)</sup> | 474.6<sup>(±5.5)</sup> | 38.1<sup>(±2.0)</sup> |
| 150 | 455.5<sup>(±3.4)</sup> | 422.6<sup>(±0.9)</sup> | 41.2<sup>(±1.1)</sup> |
| 175 | 424.4<sup>(±2.5)</sup> | 453.3<sup>(±4.1)</sup> | 39.2<sup>(±0.8)</sup> |
| 200 | 393.6<sup>(±2.7)</sup> | 465.3<sup>(±5.4)</sup> | 39.9<sup>(±1.0)</sup> |
| 225 | 380.1<sup>(±2.7)</sup> | 450.9<sup>(±4.6)</sup> | 32.8<sup>(±0.6)</sup> |

**Table 6:** ℰ and reconstruction error increase with reduced feature basis.

| *K* | ℰ (×10<sup>−4</sup>) | Reconstruction error |
|---|---|---|
| 10 | 1.6954<sup>(±0.0431)</sup> | 0.5242 |
| 25 | 1.6794<sup>(±0.0367)</sup> | 0.3596 |
| 50 | 0.8411<sup>(±0.0125)</sup> | 0.2351 |
| 75 | 0.2842<sup>(±0.0039)</sup> | 0.1358 |
| 100 | 0.0396<sup>(±0.0010)</sup> | 0.0321 |

### B.3 Running DFR-TD with a reduced feature space

Throughout our simulations we assume that the feature space of the model is the full eigenbasis where *K* = *N*, but we were curious: how robust is the model to an incomplete basis? To test this we trained and evaluated DFR-TD on the open field barrier tasks using only the top-*K* eigenvectors of *D*, sweeping *K* ∈[50, 75, 100, 125, 150, 175, 200, 225]. The main results are reported in Figure 4, here we report the mean final performance of both the DFR-MB and DFR-TD models with a reduced feature space.

### B.4 Comparison with low-rank successor representations

In the comparisons with the SR methods in the main paper we assume all of the models have the optimal sufficient rank representation for the environment or feature space. While we argue that the low-rank structure our model exploits lies in how the change is represented (more on this in Section 3.4 and Appendix B.7), comparisons to low-rank feature parameterizations are necessary to show that our method, which represents the feature space completely but the change at low rank, is better than alternative low-rank methods of representing the cognitive map.

To get at this we implement two more baseline comparisons: The first is SR-SF-*k*. We take the top-*k* eigenvectors of the base map, equivalently the eigenvectors of the random-walk SR, which is the proto-value function (PVF) / spectral-SF basis, and learn Ψ ∈ R^*N ×k*^ by ordinary TD on the ≤*k*-dimensional Bellman equation, so the induced cognitive map itself is rank *k* rather than exact. This is the standard low-rank successor implementation. The second is SR-LR (successor representation with a jointly learned low-rank factorization). We model the map directly as a product of two learned *N* ×*k* factors, Ψ ≈*LR*^*⊤*^, updating both by TD from the same sampled transitions, which is the tabular form of a forward-backward representation. Unlike SF on a frozen basis, the subspace itself adapts to the barrier rather than only the weights on fixed features. The model is also pre-trained for 50k steps in the base environment before the barrier is inserted to encourage mixing between the features.

We initialize the feature spaces of both to the top rank-*k* representation of the pre-change map. Both are given the same budget, planner temperature, step cap, and evaluation pairs as DFR-TD. Episodes are truncated at 1k steps and truncated episodes are counted at the horizon. We test the models in the same open field environment with a horizontal barrier insertion as in Section 3.1.

Neither low-rank family approaches DFR-TD at any *k*. The two behave differently as *k* grows, and neither in a way that favors compression. SR-SF-*k* improves monotonically toward *k* = *N*, so its compact regime is worse than its uncompressed one throughout. SR-LR plateaus from *k* = 50 onward, with no systematic trend thereafter; its advantage over SR-SF-*k* at small *k* reflects its ability to adapt the subspace, but this does not translate into competitive planning at any *k* we tested.

Our results establish that the correction to a predictive map has low-rank structure that the DR factorization exposes, and that this is what produces the sample-efficiency gain relative to the successor-representation baselines we test, now including genuinely low-rank ones. They do not establish that this is the only way to obtain a compact, adaptable map.

### B.5 Observable signal that the basis is insufficient under truncation

At full rank, or when the feature space captures the majority of variance of the decomposition, *F* is sufficient by construction. However, in the more relevant truncated setting *K < N*, or when the feature space does not capture all the variance, what observable signal could an agent use to detect that the current basis is insufficient?

The framework supplies an internally observable signal. Learning uses the sampled TD error from Equation (11), *δ* = *γ*(*r*_*s*_ + *f*_*s*_^*′*^ *M*) *f*_*s*_*M*. We take the same quantity per state, with the expectation over successors resolved:

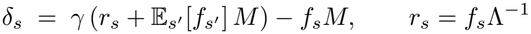

and average its squared norm over the states the agent actually visits,

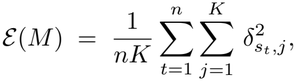

where *s*_1_, …, *s*_*n*_ are the states along a rollout of the default policy in the changed environment. The visitation frequencies are therefore supplied by the behavior itself: ℰ uses only the agent’s own operator and the transitions it experiences, and it never references the post-change map. The factor of 1*/K* makes the measure comparable across basis sizes, since *δ*_*s*_ gains a coordinate for every feature added.

We measured ℰ in the environment of Appendix B.1. For each *K* we train *M* under a fixed budget matched across conditions (400k steps, learning rate 0.001) and then evaluate ℰ once on the trained operator, from a rollout of *n* = 1000 default-policy steps taken after a 500-step burn-in. Alongside it we report the reconstruction error ∥*FMF* ^*⊤*^− *D*_new_∥ _*F*_ */*∥ *D*_new_ ∥_*F*_, the relative Frobenius error of the reconstructed map against the true post-change DR. That quantity requires knowledge of the changed environment and so is available to us but not to the agent; we include it only as an external check on the measure. Both are averaged over 20 seeds.

The measure decreases monotonically with *K*, and with the single exception of *K* = 10 versus *K* = 25, where the two values are indistinguishable, each adjacent pair is separated by more than 20 SEM across seeds. It also tracks the reconstruction error: the two agree on the ordering of the bases, fall together over the range, and reach their smallest values at *K* = *N* = 100, where the basis is complete and the residual error reflects finite training rather than truncation. Since ℰ is computed entirely from quantities internal to the agent while the reconstruction error is not, this agreement is what makes the former usable as a stand-in for the latter. We therefore propose ℰ as an intrinsic signal the agent can monitor to assess whether its current basis is adequate.

#### Choosing and expanding *K*

*K* can be grown online without discarding what has been learned. Appending an eigenvector *f*_*K*+1_ to *F* and padding *M* with a corresponding identity entry,

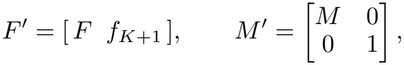

retains the existing correction and introduces the new direction at its uncorrected value, so *FMF* ^*⊤*^ gains the new spectral component and learning resumes in the enlarged basis from a warm start rather than from scratch.

The signal ℰ supplies the trigger. ℰ falls with *K* and tracks the reconstruction error the agent cannot see, so a residual that stops decreasing while learning is still in progress indicates that the basis, rather than the amount of experience, is the binding constraint. Together these give the shape of an adaptive scheme: grow *K* when the residual saturates while learning continues.

On choosing *K* in the first place, we note that in our main experiments *K* = *N* : *F* is the complete eigenbasis of the base DR, computed once and reused across environments. Both questions therefore concern the truncated regime rather than the setting our main results are reported in. Section 3.5 reports how the TD rule behaves under truncation, which is a robustness result rather than a recommendation for setting *K*. Within that regime the residual ℰ is the criterion we would propose, since it is computed from the agent’s own experience and requires no comparison against a map the agent is trying to learn.

### B.6 Composing learning of two barriers for single-shot planning

Here we show the environment (Figure 7) and the mean final performance of each model during the zero-shot composition of learned corrections (Table 7). We can see that the composed operator (*M*_1_ + *M*_2_ −*I*) achieves superior planning performance, nearing that of the model-based oracle. Using either the *M*_1_ or *M*_2_ operators alone does not provide adequate planning performance.

**Table 7:** Zero-shot composition of learned corrections Planning operator Steps to goal.

| Planning operator | Steps to goal |
|---|---|
| Open-field (no correction) | 446.9<sup>(±78.3)</sup> |
| *M*<sub>1</sub> only (barrier 1) | 420.9<sup>(±44.5)</sup> |
| *M*<sub>2</sub> only (barrier 2) | 89.6<sup>(±43.1)</sup> |
| Composed (*M*<sub>1</sub> + *M*<sub>2</sub> − *I*) | 34.7<sup>(±3.2)</sup> |
| Model-based oracle | 20.7<sup>(±0.4)</sup> |

### B.7 Basis comparison

We suspected that the advantage of our approach comes from the unique capability of the DR factorization to expose the low-rank change in a predictive map in a form that can be learned very quickly. We report the mean performance across the learning trajectory for each choice of the feature basis (Tables 8 and 9).

**Table 8:**
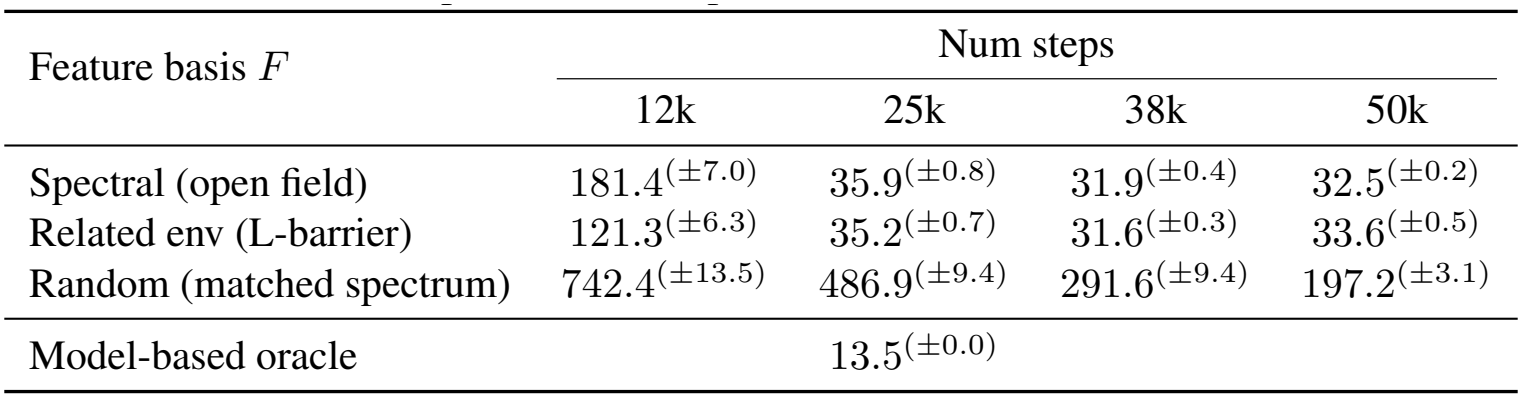
Basis comparison on the open field environment (horizontal barrier)

| Feature basis *F* | Num steps |  |  |  |
|---|---|---|---|---|
|  | 12k | 25k | 38k | 50k |
| Spectral (open field) | 181.4<sup>(±7.0)</sup> | 35.9<sup>(±0.8)</sup> | 31.9<sup>(±0.4)</sup> | 32.5<sup>(±0.2)</sup> |
| Related env (L-barrier) | 121.3<sup>(±6.3)</sup> | 35.2<sup>(±0.7)</sup> | 31.6<sup>(±0.3)</sup> | 33.6<sup>(±0.5)</sup> |
| Random (matched spectrum) | 742.4<sup>(±13.5)</sup> | 486.9<sup>(±9.4)</sup> | 291.6<sup>(±9.4)</sup> | 197.2<sup>(±3.1)</sup> |
| Model-based oracle |  | 13.5<sup>(±0.0)</sup> |  |  |

**Table 9:** Basis comparison on the four-room environment (barrier insertion)

| Feature basis *F* | Num steps |  |  |  |
|---|---|---|---|---|
|  | 37.5k | 75k | 112.5k | 150k |
| Spectral (open field) | 166.5<sup>(±8.6)</sup> | 84.4<sup>(±2.7)</sup> | 81.7<sup>(±1.8)</sup> | 81.1<sup>(±1.8)</sup> |
| Four-room (base-env DR) | 75.3<sup>(±1.2)</sup> | 70.9<sup>(±1.1)</sup> | 77.2<sup>(±2.1)</sup> | 80.7<sup>(±1.3)</sup> |
| Related env (other door blocked) | 77.8<sup>(±1.1)</sup> | 70.5<sup>(±1.1)</sup> | 77.1<sup>(±1.7)</sup> | 79.6<sup>(±1.3)</sup> |
| Random (matched spectrum) | 1000<sup>(±0.0)</sup> | 1000<sup>(±0.0)</sup> | 1000<sup>(±0.0)</sup> | 999.9<sup>(±0.1)</sup> |
| Model-based oracle |  | 27.1<sup>(±0.0)</sup> |  |  |

We can see that the advantage is fully present at *K* = *N*, so truncation cannot be its source. The random basis then shows that a full-rank basis is not sufficient either. Degradation appears only at the adversarial extreme, the random basis, which we include as a worst-case floor.

## C Experiment details

### Compute resources

All experiments were run on a MacBook Pro M2 with 16 GB of RAM running macOS Tahoe 26.1. Models and environments were implemented in custom Python code (version 3.12.12). Code for all the experiments can be found at https://github.com/arminbazza/Default-Features.

### Fixed parameters

Unless otherwise noted, we fixed the learning rate to 0.01 and the inverse temperature to 0.2 across all models and simulations. For the DFR models, the per-step cost was set to 0.1 and the terminal cost to 0. For the SR and Dyna-Q models, the per-step reward was set to 0 and the terminal reward to 10. For the reconstruction task and the simulations replicating [27], we instead used a learning rate of 0.001. We evaluate across 5 different seeds for every simulation.

### Baselines and evaluation

All three learning rules (DFR-TD, SR-TD, SR-SF) sample a state *s* uniformly from the valid set and a successor *s*^*′*^, with the same learning rate, discount, and sample budget. We write Ψ for the successor object to avoid collision with DFR’s operator *M*. As implemented, SR-SF receives the same features as DFR-TD but learns a state-indexed object, which is the relevant difference between the two. For DFR, the value function is obtained from *FMF* ^*⊤*^ via the single matrix-vector product of the linear-RL framework [23]; for the SR baselines, it is the product of the learned representation (matrix or feature-based) with the reward vector. All models act with a softmax policy (not greedy) at inverse temperature 0.2, terminating at the goal or after 1k steps.

### Open-field environment

For the open-field experiments (Sections 3.1, 3.4, & 3.5; Appendices B.3,& B.4), we considered a 15 ×15 maze under two barrier configurations. In the horizontal-barrier configuration, the barrier spanned 9 cells. In the L-shaped configuration, each arm was 7 cells long, yielding a barrier of 14 cells in total. For both configurations, we manually designated “start” and “goal” regions on opposing sides of the barrier: 3× 5 regions in the horizontal-barrier environment and 4 ×4 regions in the L-shaped environment. We then sampled 200 (start, goal) pairs uniformly from these regions, holding the same 200 pairs fixed across all evaluations. Models were trained for 50k steps with evaluation every 1k steps, and episodes were truncated at a horizon of 1k steps. For the Dyna-Q simulations, we instead fixed a single goal state and sampled 200 random starting states.

### Four-room environment

For the four-room experiments (Sections 3.2 & 3.4), we used an 11× 11 gridworld in which the walls separating rooms consisted of fully inaccessible cells, with each doorway exactly one cell wide. The agent’s starting state was fixed at the top-left corner in both the goal-revaluation and transition-revaluation settings. In the goal-revaluation condition, the goal was first placed in the top-right corner and the agent learned a representation under this configuration; the goal was then relocated to one of the eight remaining corner states and the agent was retrained. In the transition-revaluation condition, the agent was first trained with the goal in either the top-right or bottom-left room; the doorway leading to that room was then blocked and the agent was retrained. Models were trained for 20k steps with evaluation every 500 steps in the goal-revaluation setting, and for 150k steps with evaluation every 2.5k steps in the transition-revaluation setting. Each evaluation step consisted of 20 rollouts with different random seeds, with episodes truncated at 1k steps.

### Wernle merging simulation

To simulate the experiment of [27] (Section 3.6), we used a 20 ×20 environment with a barrier inserted in the middle between rows 10 and 11, partitioning the space into two 10 ×20 compartments. Taking the eigendecomposition of the fully compartmentalized environment is unsuitable as a starting feature space because the corresponding transition matrix is block-diagonal and its eigenvectors are therefore localized to a single compartment, leaving no basis for comparing how an eigenvector reorganizes across both halves. To obtain a starting representation that spans the full environment while still respecting the structure of the post-barrier dynamics, we instead computed the eigendecomposition of the open 20 ×20 space to obtain *F*_old_, and then derived *F*_new_ analytically under the post-barrier transition matrix *T*_new_. The agent was placed in the open environment with *F* initialized to *F*_new_, and the model was trained for 500k steps. For the sliding correlation analysis, we used a box width of 3.

### Reconstruction

For the reconstruction task (Appendices B.1, B.5), we used a 10 ×10 gridworld with a horizontal barrier inserted between rows 5 and 6, blocking transitions between the two halves of the environment. The barrier left two passable cells, one at each end. We trained the DFR-TD model for 800k steps and logged the reconstruction error every 1k steps. To assess sensitivity to spectral truncation, we ran the model under five values of *K*, where *K* denotes the number of top eigenvectors retained from the eigendecomposition *F* : *K* ∈ {10, 25, 50, 75, 100}.

### Composition

For the separate learning and single-shot composition of two barriers (Section 3.3) we used a 15× 15 environment with two horizontal barriers, each spanning 8 cells. The evaluation followed the same layout as in the other open-field environments described above.

## NeurIPS Paper Checklist

1. **Claims** Question: Do the main claims made in the abstract and introduction accurately reflect the paper’s contributions and scope? Answer: [Yes] Justification: The abstract’s claims are supported by theoretical and empirical evidence (Sections 2 and 3), with proofs and derivations in Appendix A.
2. **Limitations** Question: Does the paper discuss the limitations of the work performed by the authors? Answer: [Yes] Justification: See Section 4.
3. **Theory assumptions and proofs** Question: For each theoretical result, does the paper provide the full set of assumptions and a complete (and correct) proof? Answer: [Yes] Justification: We provide complete proofs or derivations for our theoretical results in Appendix A.
4. **Experimental result reproducibility** Question: Does the paper fully disclose all the information needed to reproduce the main experimental results of the paper to the extent that it affects the main claims and/or conclusions of the paper (regardless of whether the code and data are provided or not)? Answer: [Yes] Justification: We provide all details of our algorithms and baselines for reproduction in Appendix C.
5. **Open access to data and code** Question: Does the paper provide open access to the data and code, with sufficient instructions to faithfully reproduce the main experimental results, as described in supplemental material? Answer: [Yes] Justification: Our code is released at https://github.com/arminbazza/Default-Features.
6. **Experimental setting/details** Question: Does the paper specify all the training and test details (e.g., data splits, hyperparameters, how they were chosen, type of optimizer) necessary to understand the results? Answer: [Yes] Justification: The main text provides experimental settings sufficient for understanding, with additional details, including hyperparameters in Appendix C.
7. **Experiment statistical significance** Question: Does the paper report error bars suitably and correctly defined or other appropriate information about the statistical significance of the experiments? Answer: [Yes] Justification: Figures report means with shaded 95% confidence intervals; tables report means with SEM in parentheses. Both are computed across 5 seeds.
8. **Experiments compute resources** Question: For each experiment, does the paper provide sufficient information on the computer resources (type of compute workers, memory, time of execution) needed to reproduce the experiments? Answer: [Yes] Justification: We discuss compute resources in Appendix C.
9. **Code of ethics** Question: Does the research conducted in the paper conform, in every respect, with the NeurIPS Code of Ethics https://neurips.cc/public/EthicsGuidelines? Answer: [Yes] Justification: This paper conforms with the NeurIPS Code of Ethics.
10. **Broader impacts** Question: Does the paper discuss both potential positive societal impacts and negative societal impacts of the work performed? Answer: [No] Justification: While RL broadly might be used for applications with both positive and negative outcomes, our algorithmic contributions are not tied to any particular application.
11. **Safeguards** Question: Does the paper describe safeguards that have been put in place for responsible release of data or models that have a high risk for misuse (e.g., pre-trained language models, image generators, or scraped datasets)? Answer: [No] Justification: We don’t believe our model poses any risks or obvious potentials for misuse.
12. **Licenses for existing assets** Question: Are the creators or original owners of assets (e.g., code, data, models), used in the paper, properly credited and are the license and terms of use explicitly mentioned and properly respected? Answer: [Yes] Justification: All original owners of the code used in the paper are properly credited, and license information is mentioned.
13. **New assets** Question: Are new assets introduced in the paper well documented and is the documentation provided alongside the assets? Answer: [N/A] Justification: The paper does not introduce new datasets, pre-trained models, or software libraries as released assets.
14. **Crowdsourcing and research with human subjects** Question: For crowdsourcing experiments and research with human subjects, does the paper include the full text of instructions given to participants and screenshots, if applicable, as well as details about compensation (if any)? Answer: [N/A] Justification: The paper does not involve crowdsourcing nor research with human subjects.
15. **Institutional review board (IRB) approvals or equivalent for research with human subjects** Question: Does the paper describe potential risks incurred by study participants, whether such risks were disclosed to the subjects, and whether Institutional Review Board (IRB) approvals (or an equivalent approval/review based on the requirements of your country or institution) were obtained? Answer: [N/A] Justification: The paper does not involve crowdsourcing nor research with human subjects.
16. **Declaration of LLM usage** Question: Does the paper describe the usage of LLMs if it is an important, original, or non-standard component of the core methods in this research? Note that if the LLM is used only for writing, editing, or formatting purposes and does *not* impact the core methodology, scientific rigor, or originality of the research, declaration is not required. Answer: [N/A] Justification: LLMs have been used for coding assistance, drafting, editing, and formatting purposes.

## Footnotes

1 Code: https://github.com/arminbazza/Default-Features

## Notes

### Competing Interest Statement

The authors have declared no competing interest.

### Summary of Updates

We finalized the manuscript for the camera ready version for NeurIPS

## References

[1] Edward C Tolman. Cognitive maps in rats and men. Psychological review, 55(4):189, 1948.

[2] John O’Keefe and Jonathan Dostrovsky. The hippocampus as a spatial map: preliminary evidence from unit activity in the freely-moving rat. Brain research, 1971.

[3] John O’keefe and Lynn Nadel. The hippocampus as a cognitive map. Oxford university press, 1978.

[4] Bruce L McNaughton, Francesco P Battaglia, Ole Jensen, Edvard I Moser, and May-Britt Moser. Path integration and the neural basis of the’cognitive map’. Nature Reviews Neuroscience, 7(8):663–678, 2006.

[5] Timothy EJ Behrens, Timothy H Muller, James CR Whittington, Shirley Mark, Alon B Baram, Kimberly L Stachenfeld, and Zeb Kurth-Nelson. What is a cognitive map? organizing knowledge for flexible behavior. Neuron, 100(2):490–509, 2018.

[6] James CR Whittington, Timothy H Muller, Shirley Mark, Guifen Chen, Caswell Barry, Neil Burgess, and Timothy EJ Behrens. The tolman-eichenbaum machine: unifying space and relational memory through generalization in the hippocampal formation. Cell, 183(5):1249–1263, 2020.

[7] Dileep George, Rajeev V Rikhye, Nishad Gothoskar, J Swaroop Guntupalli, Antoine Dedieu, and Miguel Lázaro-Gredilla. Clone-structured graph representations enable flexible learning and vicarious evaluation of cognitive maps. Nature communications, 12(1):2392, 2021.

[8] Peter Dayan. Improving generalization for temporal difference learning: The successor representation.Neural computation, 5(4):613–624, 1993.

[9] Kimberly L Stachenfeld, Matthew M Botvinick, and Samuel J Gershman. The hippocampus as a predictive map. Nature neuroscience, 20(11):1643–1653, 2017.

[10] Ida Momennejad, Evan M Russek, Jin H Cheong, Matthew M Botvinick, Nathaniel Douglass Daw, and Samuel J Gershman. The successor representation in human reinforcement learning. Nature human behaviour, 1(9):680–692, 2017.

[11] Evan M Russek, Ida Momennejad, Matthew M Botvinick, Samuel J Gershman, and Nathaniel D Daw. Predictive representations can link model-based reinforcement learning to model-free mechanisms. PLoS computational biology, 13(9):e1005768, 2017.

[12] Jesse P Geerts, Samuel J Gershman, Neil Burgess, and Kimberly L Stachenfeld. A probabilistic successor representation for context-dependent learning. Psychological Review, 131(2):578, 2024.

[13] Armin Bazarjani and Payam Piray. Efficient learning of predictive maps for flexible planning. bioRxiv, pages 2026–02, 2026.

[14] Armin Bazarjani, Caitlin Mallory, David Foster, and Payam Piray. Hippocampal replay repairs the cognitive map for flexible planning. bioRxiv, pages 2026–10, 2026.

[15] Tejas D Kulkarni, Ardavan Saeedi, Simanta Gautam, and Samuel J Gershman. Deep successor reinforcement learning. arXiv preprint arXiv:1606.02396, 2016.

[16] André Barreto Will Dabney, Rémi Munos, Jonathan J Hunt, Tom Schaul, Hado P Van Hasselt, and David Silver. Successor features for transfer in reinforcement learning. Advances in neural information processing systems, 30, 2017.

[17] Diana Borsa, André Barreto John Quan, Daniel Mankowitz, Rémi Munos, Hado Van Hasselt, David Silver, and Tom Schaul. Universal successor features approximators. arXiv preprint arXiv:1812.07626, 2018.

[18] Andre Barreto, Diana Borsa, John Quan, Tom Schaul, David Silver, Matteo Hessel, Daniel Mankowitz, Augustin Zidek, and Remi Munos. Transfer in deep reinforcement learning using successor features and generalised policy improvement. In International Conference on Machine Learning, pages 501–510. PMLR, 2018.

[19] Ahmed Touati and Yann Ollivier. Learning one representation to optimize all rewards. Advances in Neural Information Processing Systems, 34:13–23, 2021.

[20] Benjamin Eysenbach, Tianjun Zhang, Sergey Levine, and Russ R Salakhutdinov. Contrastive learning as goal-conditioned reinforcement learning. Advances in Neural Information Processing Systems, 35: 35603–35620, 2022.

[21] Michael Janner, Igor Mordatch, and Sergey Levine. γ-models: Generative temporal difference learning for infinite-horizon prediction. In Advances in Neural Information Processing Systems, volume 33, 2020.

[22] Lucas Lehnert and Michael L Littman. Successor features combine elements of model-free and model-based reinforcement learning. Journal of Machine Learning Research, 21(196):1–53, 2020.

[23] Payam Piray and Nathaniel D Daw. Linear reinforcement learning in planning, grid fields, and cognitive control. Nature communications, 12(1):4942, 2021.

[24] Payam Piray and Nathaniel D Daw. Reconciling flexibility and efficiency: medial entorhinal cortex represents a compositional cognitive map. Nature communications, 16(1):7444, 2025.

[25] John Tsitsiklis and Benjamin Van Roy. Analysis of temporal-diffference learning with function approximation. Advances in neural information processing systems, 9, 1996.

[26] Richard S Sutton, Andrew G Barto, et al. Reinforcement learning: An introduction, volume 1. MIT press Cambridge, 1998.

[27] Tanja Wernle, Torgeir Waaga, Maria Mørreaunet, Alessandro Treves, May-Britt Moser, and Edvard I Moser. Integration of grid maps in merged environments. Nature neuroscience, 21(1):92–101, 2018.

[28] Emanuel Todorov. Linearly-solvable markov decision problems. Advances in neural information processing systems, 19, 2006.

[29] Emanuel Todorov. Efficient computation of optimal actions. Proceedings of the national academy of sciences, 106(28):11478–11483, 2009.

[30] William W Hager. Updating the inverse of a matrix. SIAM review, 31(2):221–239, 1989.

[31] Gene H Golub and Charles F Van Loan. Matrix computations. JHU press, 2013.

[32] Richard S Sutton. Learning to predict by the methods of temporal differences. Machine learning, 3(1): 9–44, 1988.

[33] Steven J Bradtke and Andrew G Barto. Linear least-squares algorithms for temporal difference learning. Machine learning, 22(1):33–57, 1996.

[34] Sridhar Mahadevan and Mauro Maggioni. Proto-value functions: A laplacian framework for learning representation and control in markov decision processes. Journal of Machine Learning Research, 8(10), 2007.

[35] Marlos C Machado, Marc G Bellemare, and Michael Bowling. A laplacian framework for option discovery in reinforcement learning. In International conference on machine learning, pages 2295–2304. PMLR, 2017.

[36] Richard S Sutton. Integrated architectures for learning, planning, and reacting based on approximating dynamic programming. In Machine learning proceedings 1990, pages 216–224. Elsevier, 1990.

[37] Richard S Sutton. Dyna, an integrated architecture for learning, planning, and reacting. ACM Sigart Bulletin, 2(4):160–163, 1991.

[38] Dharshan Kumaran, Demis Hassabis, and James L McClelland. What learning systems do intelligent agents need? complementary learning systems theory updated. Trends in cognitive sciences, 20(7):512–534, 2016.

[39] Jane X Wang, Zeb Kurth-Nelson, Dharshan Kumaran, Dhruva Tirumala, Hubert Soyer, Joel Z Leibo, Demis Hassabis, and Matthew Botvinick. Prefrontal cortex as a meta-reinforcement learning system. Nature neuroscience, 21(6):860–868, 2018.

[40] Matthew Botvinick, Sam Ritter, Jane X Wang, Zeb Kurth-Nelson, Charles Blundell, and Demis Hassabis. Reinforcement learning, fast and slow. Trends in cognitive sciences, 23(5):408–422, 2019.

[41] Torkel Hafting, Marianne Fyhn, Sturla Molden, May-Britt Moser, and Edvard I Moser. Microstructure of a spatial map in the entorhinal cortex. Nature, 436(7052):801–806, 2005.

[42] Hanne Stensola, Tor Stensola, Trygve Solstad, Kristian Frøland, May-Britt Moser, and Edvard I Moser. The entorhinal grid map is discretized. Nature, 492(7427):72–78, 2012.

[43] Marianne Fyhn, Torkel Hafting, Alessandro Treves, May-Britt Moser, and Edvard I Moser. Hippocampal remapping and grid realignment in entorhinal cortex. Nature, 446(7132):190–194, 2007.

[44] Stefan Leutgeb, Jill K Leutgeb, Carol A Barnes, Edvard I Moser, Bruce L McNaughton, and May-Britt Moser. Independent codes for spatial and episodic memory in hippocampal neuronal ensembles. Science, 309(5734):619–623, 2005.

[45] Laura Lee Colgin, Edvard I Moser, and May-Britt Moser. Understanding memory through hippocampal remapping. Trends in neurosciences, 31(9):469–477, 2008.

[46] Øyvind Arne Høydal, Emilie Ranheim Skytøen, Sebastian Ola Andersson, May-Britt Moser, and Edvard I Moser. Object-vector coding in the medial entorhinal cortex. Nature, 568(7752):400–404, 2019.

[47] Dori Derdikman, Jonathan R Whitlock, Albert Tsao, Marianne Fyhn, Torkel Hafting, May-Britt Moser, and Edvard I Moser. Fragmentation of grid cell maps in a multi compartment environment. Nature neuroscience, 12(10):1325–1332, 2009.

[48] Nathaniel D Daw, Yael Niv, and Peter Dayan. Uncertainty-based competition between prefrontal and dorsolateral striatal systems for behavioral control. Nature neuroscience, 8(12):1704–1711, 2005.

